# Two hundred years of changes in orchid pollination revealed using herbarium specimens

**DOI:** 10.64898/2025.12.05.692704

**Authors:** Lola Torres-Montagner, André Schuiteman, Joanne M Bennett, Tiffany Knight, Demetra Rakosy, Michael F. Fay, Philip C. Stevenson, Carlos Martel

## Abstract

Orchids are one of the most diverse plant families and are renowned for their highly specialised interactions with pollinators. Despite growing evidence of orchid population decline, pollinator decline and rising concerns for orchid reproductive success, long-term changes remain poorly understood. To address this gap, we analysed changes in pollinarium removal, as a proxy for pollination success, in herbarium specimens of three species-rich orchid genera (*Disa, Oncidium* and *Ophrys*) hosted at the Royal Botanic Gardens, Kew, which has a rich and diverse orchid collection spanning the past two centuries. The selected genera occur in different major regions (subtropical and tropical Africa, predominantly temperate Europe, and tropical America, respectively). Our analysis reveals that pollinarium removal declined significantly in *Disa* and *Oncidium*, particularly among species with deceptive strategies or specialised pollination mechanisms. In contrast, *Ophrys* showed a significant increase, driven by Apidae-pollinated species, whereas pollinarium removal for Andrenidae-pollinated species declined over time. These contrasting findings reflect the role of orchid identity, pollinator availability and pollination strategy in shaping reproductive success under anthropogenic change. Overall, this study demonstrates the power of herbarium specimens to reveal long-term ecological changes, providing unique insights into the response of specific plant-pollinator interactions to increasing anthropogenic pressure.

## Introduction

Pollination and pollinators are essential for sustaining biodiversity, maintaining ecosystem health, supporting key ecosystem services, and ensuring global food security. Roughly 80% of plants are dependent on animal pollinators for reproduction (Rodger et al. 2021). Nevertheless, the largely mutualistic and intricate interactions between plants and their pollinators are increasingly threatened by anthropogenic pressures that lead to habitat fragmentation, widespread declines in both plant and pollinator populations, and the overall health of natural ecosystems (Augspurger, 2017; Biesmeijer et al., 2006; Hailay Gebremariam, 2024; Raven and Wagner, 2021; Vellend et al., 2017). Effectively, recent studies have shown that environmental and anthropogenic changes disrupt predominantly specialised plant-pollinator interactions (Zoller et al. 2023), such as interactions between some orchid species with restricted geographic range and their specialised pollinators, which are therefore likely to be lost.

The orchid family (Orchidaceae) is a cosmopolitan and extremely diverse plant group, comprising over 28,000 species and 736 recognised genera (Chase et al., 2015). Unfortunately, orchids are one of the most at-risk plant families in the world (Fay et al. 2025) and are particularly susceptible to anthropogenic activities and climate change. Currently, more than half of the assessed orchid species are listed in one of the categories of threat according to the IUCN Global Red List (Gale et al., 2018; Fay et al., 2025), and they are also the most pollen-limited plant family (Bennett et al., 2020). Environmental changes have led to changes in phenology in orchid species, resulting in desynchronization with pollinator activity and therefore affecting pollinator availability and orchid pollination success (Schiestl et al., 2025). Orchids are known for their intricate relationships with pollinators and attractive flowers, which have evolved various pollination mechanisms (e.g. generalised food deception, sexual deception) (Jersáková et al., 2006; Capó et al., 2023), and have adapted to diverse pollinators (e.g. bees, beetles, birds, butterflies, flies, moths; Ackerman et al., 2023). Orchids and their pollination have long attracted early botanists and naturalists, including Charles Darwin, whose work “On the various contrivances by which British and foreign orchids are fertilised by insects” was used to illustrate how floral adaptations are a consequence of natural selection (Darwin, 1862, 1877). To date, orchids are among the most extensively studied plant families in terms of pollination, even though the pollination mechanisms of only about 5% of their species are known (Ackerman et al., 2023).

For centuries, plants have been collected for ornamental purposes, although it was during the Renaissance that plant collecting for botanical gardens in universities began. However, the systematic, large-scale collection of plants for economic and scientific purposes by botanical gardens expanded significantly with the rise of colonial empires and the scientific expeditions of the 18th and 19th centuries. Explorers and plant collectors travelled to natural areas around the world to discover and bring new orchids and other plants to enrich private collections and botanical gardens, many of which also ended up in herbaria. Herbarium specimens offer a unique window into the past, allowing for long-term studies on different biological and ecological aspects, including pollination interactions (Eckert et al., 2025), such as the evaluation of temporal changes in plant-pollinator networks (Rakosy et al., 2023; Daru and Zhigila, 2025). The study of changes in pollination rates using herbarium specimens can serve as a powerful proxy for biodiversity loss, since changes in ecosystems and the availability of pollinators or shifts in flowering patterns will affect plant reproductive success over time (Rodger et al., 2021). While the impact of anthropogenic change on pollinator and plant populations has been extensively studied on a short time scale, fewer studies have been investigating temporal patterns in reproductive success using long-term data sets (e.g. Song et al. 2025), which combine data from historical collections and re-collected contemporary specimens (Burkle et al., 2013; Zoller et al., 2023). However, most of the existing literature on plant-pollinator interactions focuses on a limited number of species within the predominantly temperate regions of Europe, North America and Australia, leaving the highly diverse tropics overlooked and their specimens underutilised (Park et al., 2023).

Orchids can be used as a model system to investigate changes in pollination rates because they are widely represented in major herbaria. Unlike other plant groups, many orchid species bear large flowers, and all have evolved a pollinarium, a composite structure that includes two or more pollinia, which are coherent masses of pollen grains, and associated accessory structures. The latter structures facilitate attachment to the adapted pollinator, and therefore, the presence or absence of pollinarium is strongly linked to pollinator availability and pollination success. Although pollination rate in orchid specimens can be studied by rehydrating and dissecting flowers to then look at pollinarium removal (Pauw and Hawkins 2010), the occurrence of the pollinarium allows the identification of the presence or absence of pollinaria on orchids using magnifying lenses or stereomicroscopes, without compromising the integrity of herbarium specimens. In this study, we aimed to identify the changes in pollination rates over time in three orchid genera, each from a different continent, in relation to the pollination mechanism, geographical area and pollinator type.

## Materials and Methods

### Plant genera selection

The criteria for the selection of the sample genera were the sample size (high number of specimens), genus richness (at least 20 species in the genus), their distribution (wide distribution), and the presence of different pollination mechanisms and life forms. The general pressing method for each genus had to allow for the assessment of pollinarium removal, meaning that the anther cap had to be visible. *Disa, Oncidium*, and *Ophrys* fitted the criteria and were selected for this study.

*Disa*: this genus includes 185 accepted species of terrestrial orchids native to tropical South Africa (POWO, 2025). We selected the section *Micranthae*, as it is the most diverse and is well represented in Kew’s collection. The known pollination mechanisms are butterfly pollination, moth pollination, bird pollination, fly pollination and bee pollination (Byers, 2021; Johnson et al., 1998a). Both rewarding and deceptive strategies occur in the genus (Johnson et al., 2013, 1998b).

*Oncidium*: this genus includes 335 accepted species of epiphytic orchids native to tropical and subtropical America (POWO, 2025). Most *Oncidium* species rely on oil-collecting bees for pollination. Both rewarding and deceptive strategies occur in the genus (Pansarin et al., 2017; Parra-Tabla et al., 2000).

*Ophrys*: this genus includes 25 accepted species of terrestrial orchids native to Macaronesia, Europe to Caucasus, the Mediterranean basin to South Turkmenistan (POWO, 2025). *Ophrys* flowers are renowned for their specialisation in sexual deception and specific interactions with pollinators (Schatz et al., 2020). They do not offer any reward to pollinators (Jersáková et al., 2006).

### Herbarium sampling

Orchid specimens deposited at the herbarium of the Royal Botanic Gardens, Kew (K), one of the largest herbaria in the world, were inspected using a stereomicroscope. For terrestrials like *Ophrys* and *Disa*, one herbarium specimen could contain more than one individual. In this case, when the number of plants on the specimen was even, the first one on the left was selected. On a specimen with an uneven number of individuals, the middle plant was selected. For each specimen, the collection year, the species name, and the location were collected. The total number of opened flowers, with the anther cap visible, was counted. Each flower was assessed for pollinarium removal. The absence of one or both pollinia indicated pollinarium removal, and the flower was then considered to have successfully exported its pollinia by pollinator means (male success). This was then reported as “number of pollinaria removed” by flower and divided by the total number of flowers to obtain a ratio per specimen. However, when the deposited pollinia were still attached to the caudicle or stipe, they were classified as self-pollination. When the assessment of pollinarium removal was not possible due to bad conditions of the specimen (e.g. mould, pre-pressing damage), the absence of flowers on the specimens, or pressing of the flower facing down, the specimens were not included in the posterior statistical analyses. The method for assessing pollinarium removal had to be consistent throughout, but the criterion chosen to decide if pollination had occurred naturally was specific to each genus. For species with loose pollinia, like in *Ophrys* and *Disa*, the pollinia were sometimes damaged, but some of them were still in the anther cap or visible on a lower flower. In this case, removal was most likely artificial during or after the specimen collection and, therefore, were specimens not included in the posterior analyses. For *Oncidium*, if the pollinia were not present but the stipe was still visible under or next to the anther cap, removal was considered artificial and not accounted for. Species with a predominantly spontaneous self-pollination strategy (e.g. *Ophrys apifera*) were not included in the analysis.

### Statistical analysis

We analysed the probability of pollinarium removal as a function of collection year, pollinator type, and geographical region using generalised linear models (GLMs) and generalised linear mixed-effects models (GLMMs), both with binomial distributions and logit link functions. The collection year was scaled to mean zero and unit variance prior to analysis. To account for possible non-independence of samples from the same region, we also fitted GLMMs with geographical region as a random effect. The locations were grouped into regions to improve the model fit. A subset of the data containing only the species for which we had information on the pollinator identity was created, where pollinator type was a categorical variable (Supplementary Table 1). Pollinator information was extracted from Ackerman et al. (2023) for *Disa* and *Oncidium* and from Claessens and Kleynen (2016) for *Ophrys*. When no pollinator information was available, the pollinator type was inferred based on the species’ features, or the species was not included in the analysis. GLMs were then fitted to the subset by modelling the number of pollinaria removed per specimen as a function of scaled collection date, pollinator group, and their interaction. We conducted Bayesian changepoint analysis at a threshold of 0.01 to detect significant shifts in the mean probability of pollinarium removal. GLMs and GLMMs analyses were conducted using the lme4 package (Bates et al., 2015), shifts over time in pollinarium removal using the Rbeast package (Zhao et al., 2019) and data visualisation was performed using the Tidyverse package (Wickham et al., 2019), all in the R software.

## Results

In total,1809 orchid specimens were examined, representing 114 species within the three genera. In the following section, pollination rate patterns are presented for each genus.

### Disa

Pollinarium removal significantly decreased over time (GLM, χ^2^ = 91.29, df = 1, p < 0.001) for *Disa* specimens. However, no interaction effect of collection date and regions on pollinarium removal was found (GLM, χ^2^ = 2.90, df = 2, p = 0.235). While South Africa showed a marginally significant lower baseline probability (p = 0.05), temporal trends were stable and similar across tropical Africa (Figure 1). In the subset of 89 specimens with available pollinator information, pollinarium removal over time significantly varied among pollinator groups (GLM, χ^2^ =28.62, df = 3, p < 0.001). Species pollinated by a wide variety of pollinators, like the generalist *Disa fragrans*, and species pollinated by Nectariniidae and Sphingidae had a low initial pollinarium removal probability but declined at slow rate. However, *Disa* species pollinated by Scoliidae exhibited a steep decline over time, significantly different from the other groups (p = 0.0051; Figure 2). Bayesian changepoint analysis identified four significant shifts (threshold = 0.01). There was an abrupt decrease in pollinarium removal over time in 1924 (change = - 0.022), 1969 (change = - 0.025) and 1981 (change = - 0.011).

**Figure 1.**
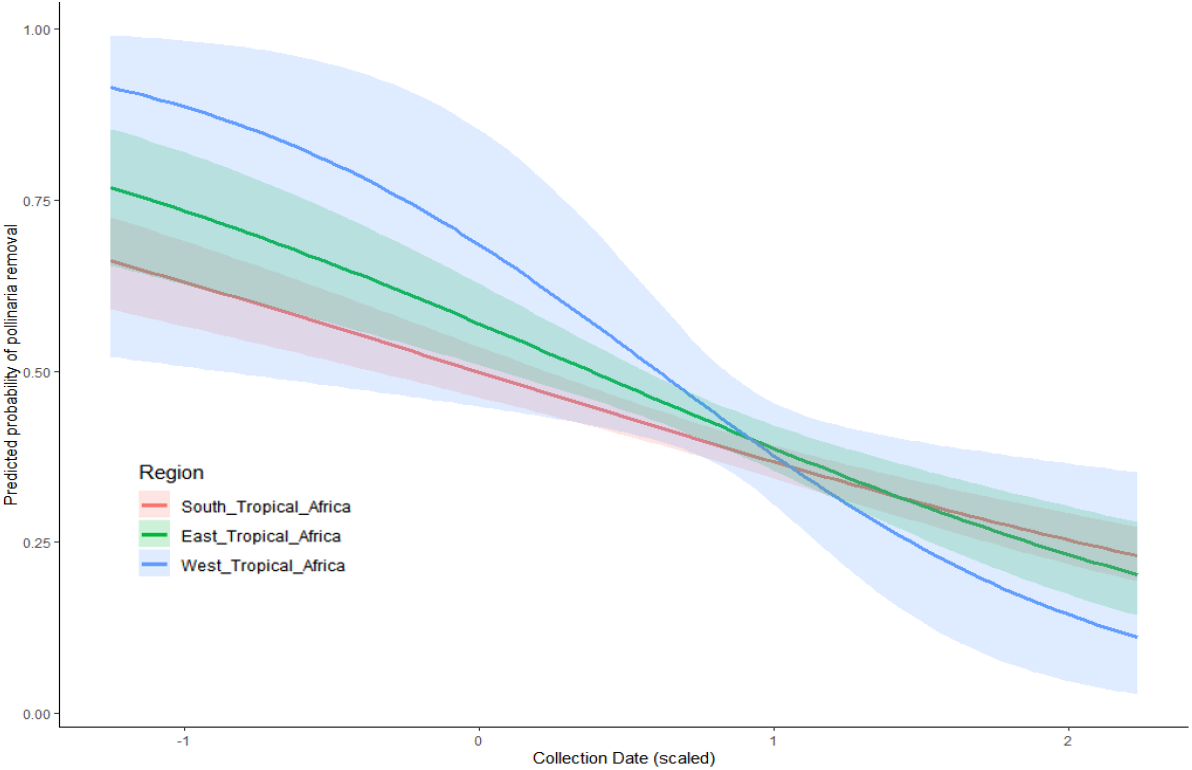
Predicted probability of pollinarium removal in *Disa* over time across three African regions. Shaded areas indicate 95% confidence intervals. All regions show a significant decline over time in pollinarium removal probability, with South Tropical Africa maintaining a lower baseline.

**Figure 2.**
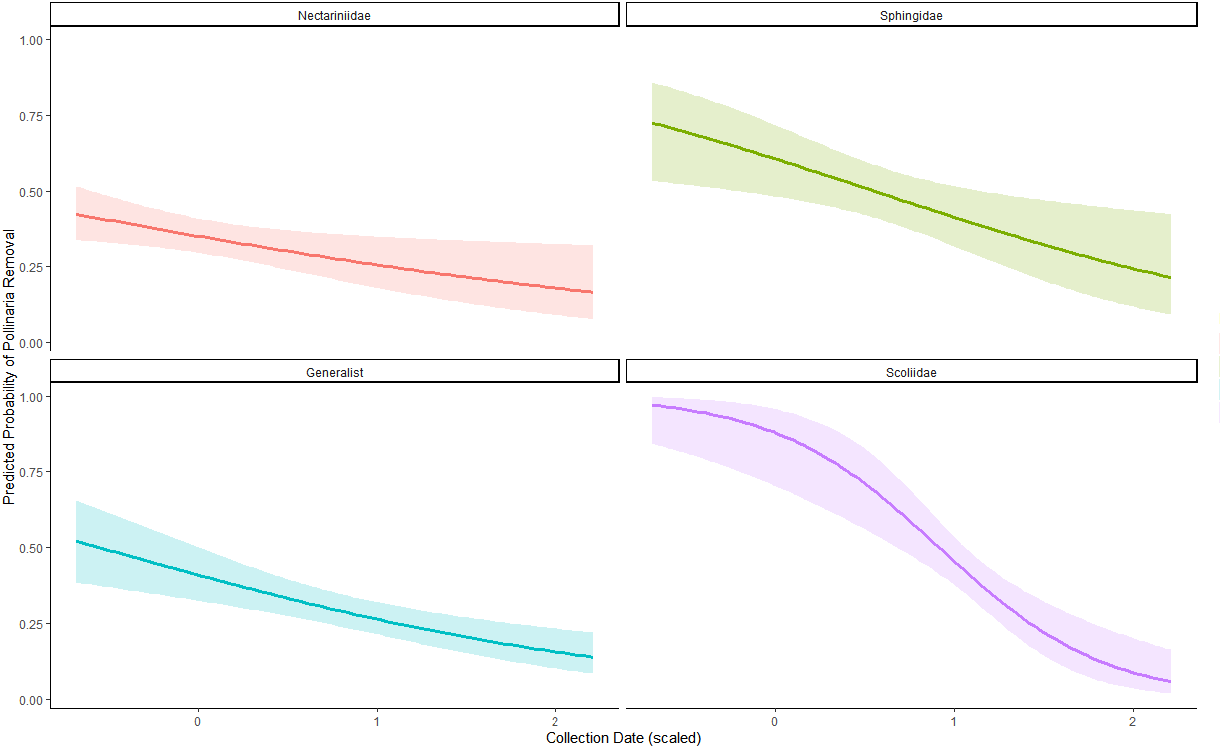
Predicted probability of pollinarium removal in *Disa* over time by pollinator group. Results are from a subset of species with known pollinators (n = 89). Shaded ribbons represent 95% confidence intervals. A significant interaction was detected between pollinator group and collection date.

### Oncidium

Pollinarium removal probability significantly declined over time (GLM, χ^2^ =186.32, df = 1, p < 0.001). No significant interaction was found between collection date and region (GLM, χ^2^ = 2.63, df = 3, p = 0.45), indicating that pollinarium removal declined across South America (Figure 3). The analysis on the subset of specimens with strategy information revealed that pollinarium removal over time significantly differs between “Deceit” and “Reward” strategies (GLM, χ^2^ = 5.42, df = 1, p < 0.05). Specimens associated with deceptive strategies showed a sharp decline in removal probability over time. In contrast, rewarding species showed a significantly different (p = 0.019), more stable trend (Figure 4). The Bayesian changepoint analysis did not reveal any changepoint above the threshold for *Oncidium*, which indicates the pollination rate has decreased constantly in the last two hundred years.

**Figure 3.**
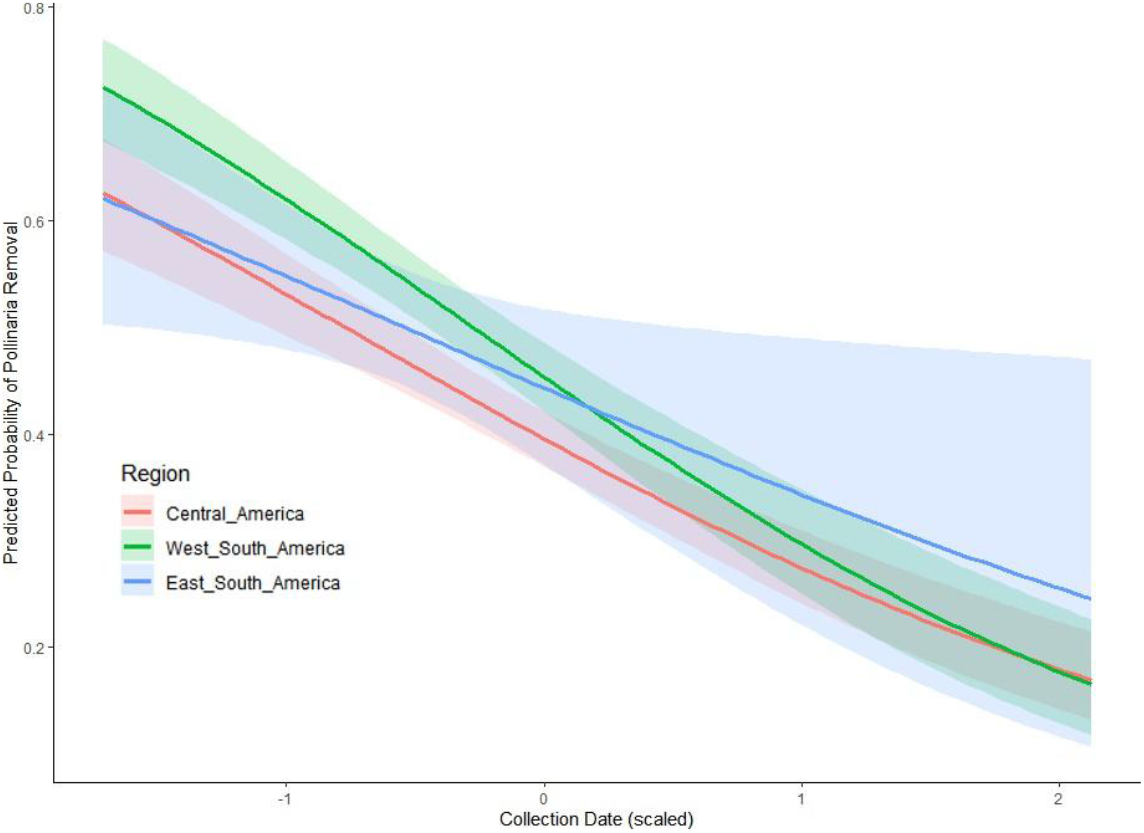
Predicted probability of pollinarium removal in *Oncidium* over time across three Neotropical regions. Shaded areas indicate 95% confidence intervals. All regions show a significant decline over time.

**Figure 4.**
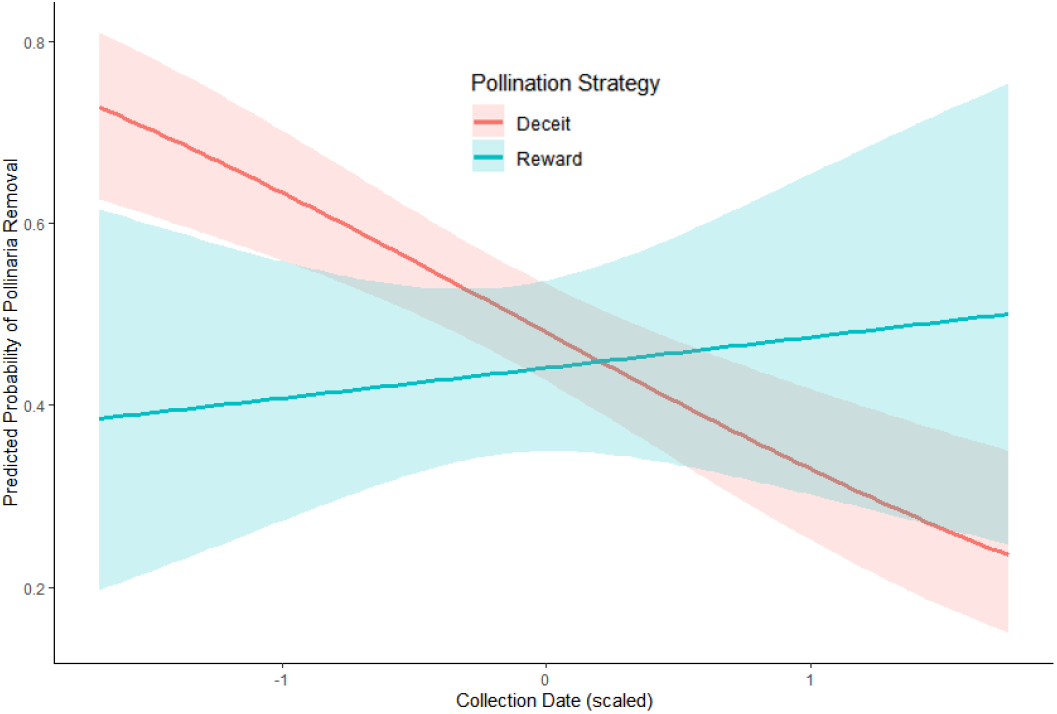
Predicted probability of pollinarium removal in *Oncidium* over time by pollinator strategy. Model predictions are from a subset of 48 specimens with known strategy. Shaded areas indicate 95% confidence intervals. Deceptive species show a sharp decline over time, while rewarding species remain more stable.

### Ophrys

Pollinarium removal probability over time in *Ophrys* species significantly increased over time (GLM, χ^2^ = 5.6201, df = 1, p < 0.05). No significant effects of region were detected; the positive trend persisted across Europe. Pollinator family significantly influenced trends in pollinarium removal over time (GLM, χ^2^ = 8.1069, df = 3, p < 0.05) as seen in Figure 5. For Andrenidae pollinated *Ophrys*, there was a marginally significant trend towards a decrease in pollinarium removal over time (p = 0.057). In contrast, there was a significant increase in pollinarium removal for Apidae pollinated *Ophrys* (p = 0.007). The Bayesian changepoint analysis did not reveal any changepoint above the threshold for *Ophrys*, which indicates the pollination rate has steadily increased in the last two hundred years.

**Figure 5.**
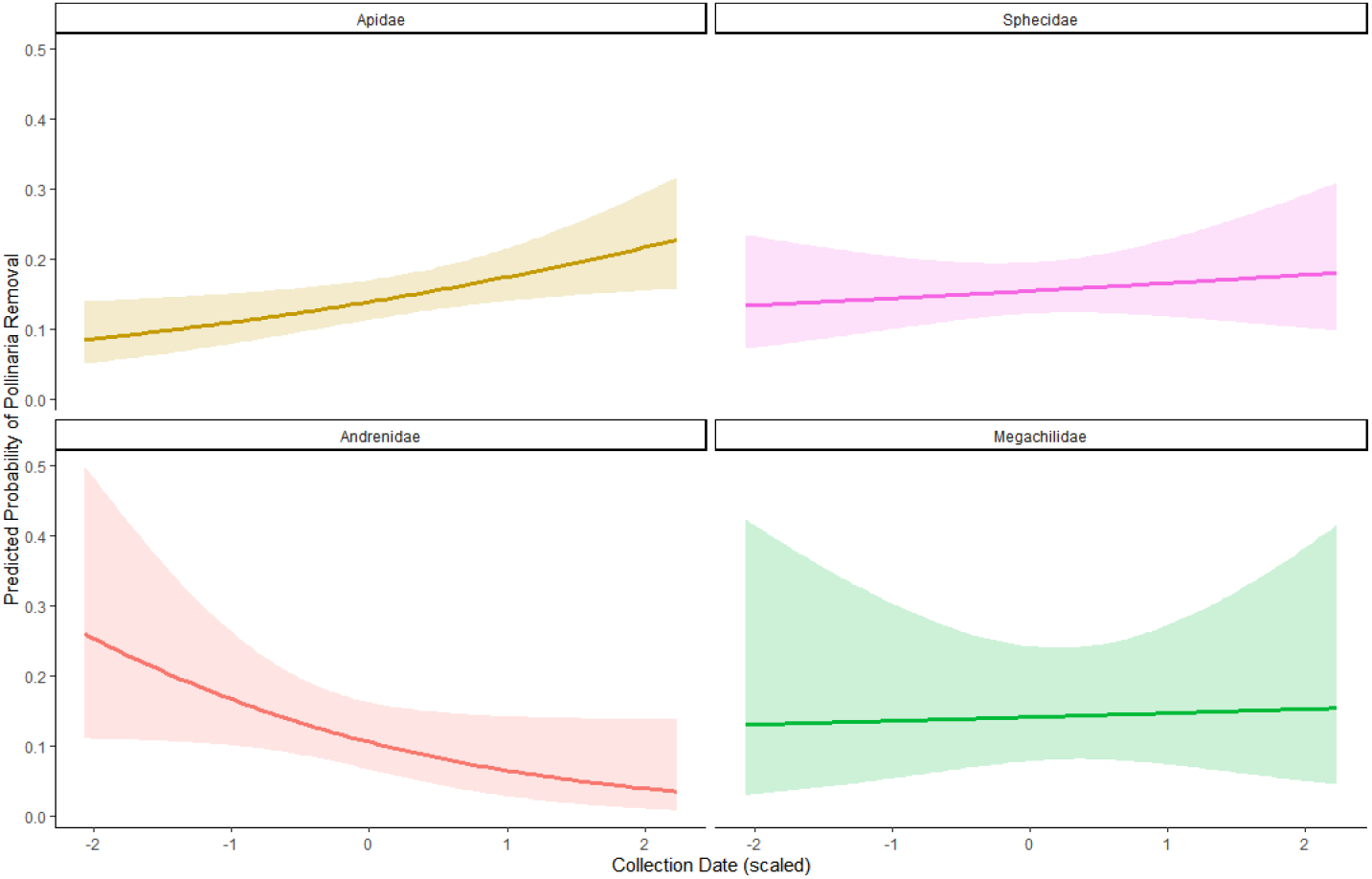
Predicted probability of pollinarium removal in *Ophrys* over time by pollinator family. Shaded areas indicate 95% confidence intervals. Pollinarium removal increased over time in Apidae-pollinated species and declined in Andrenidae-pollinated species, while no significant changes were detected for Megachilidae and Sphecidae.

### Overall trends

Pollinarium removal significantly changed over the last 200 years and was affected by the genus identity (Figure 6). Pollinarium removal significantly declined over time for the reference genus *Disa* (GLM, χ^2^ = 82053, df = 2, p < 0.001), and the trend for *Oncidium* did not differ from it (p = 0.87). The trend for *Ophrys* significantly differed from *Disa* and *Oncidium* (p < 0.0001).

**Figure 6.**
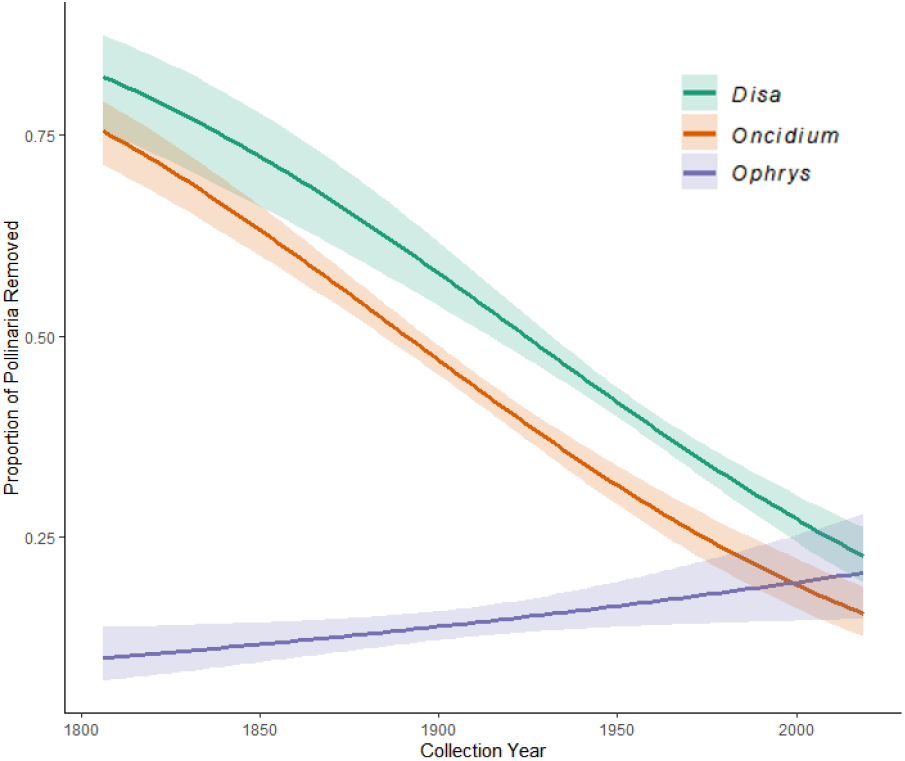
Predicted probability of pollinarium removal over time in *Disa, Oncidium* and *Ophrys*. Shaded areas indicate 95% confidence intervals. *Disa* and *Oncidium* show significant declines, whereas *Ophrys* shows a significant increase.

## Discussion

This study represents a significant advancement in utilising herbarium specimens for ecological studies, as we uncovered changes in pollination rates in orchid genera, reflecting reproductive trends in understudied and threatened pollinator species. Across the three genera, pollinarium removal significantly varied, reflecting differences in habitats, pollinator identity and pollination strategy. Both *Disa* and *Oncidium* exhibited significant declines in pollinarium removal over time, consistent with the widespread loss of pollinators in Africa and South America and highlighting the vulnerability of specific interactions in tropical environments. In contrast, *Ophrys* showed an increase in removal rates across Europe, suggesting that some European pollinators exhibit a greater tolerance to anthropogenic pressures or an increased efficiency in pollination due to yet unknown factors. This resilience is possibly linked to the fact that plant-pollinator interactions and the impact of biodiversity loss have been extensively studied in temperate regions of Europe, and protected areas have already been put in place. In contrast, natural habitats in Africa and South America are facing intense pressure from activities such as mining and agricultural expansion (Laurance et al., 2014). In the following sections, we will discuss the results for each genus in more detail.

### Disa

In *Disa*, we observed a significant long-term decline in pollinarium removal over time across tropical South Africa, indicating a decrease in visitation and reproduction success across the section *Micranthae. Disa* species are highly specialised and are therefore especially sensitive to pollinator decline and habitat loss (Johnson, 1995; Johnson et al., 1998a, 2024). These findings support concerns over the decline of insects in African tropical regions, as shown in a global analysis on the effect of land use on pollinator diversity, where pollinator abundance was found to be decreasing by 75% in intensive pastures (Millard et al., 2023). Our study also identified habitat loss and climate change as key drivers of pollinator decline in tropical regions (Millard et al., 2021). A report from the Intergovernmental Science-Policy Platform on Biodiversity and Ecosystem Services (IPBES) stresses that agricultural intensification, mining, use of pesticides and habitat fragmentation in South Africa are tightly linked to the decline of pollinators (Archer et al., 2018). The results from the Bayesian change point analysis could reflect the intensive agricultural expansion and pesticide use during this period.

The analysis of pollinator types revealed that trends in pollinarium removal varied depending on the pollinator identity. Particularly, species pollinated by Scoliidae exhibited a steep and significantly sharper decline in removal probability over time compared to species associated with Sphingidae, Nectariniidae and generalists that showed a more gradual decline. These findings potentially reflect the sensitivity of specialised species to anthropogenic changes, as Scoliidae are specialised parasitic wasps that rely on their host for survival. Recent studies show that specialised reproductive interactions are more threatened by climate change and anthropogenic activity than generalist interactions, as they can adapt to a wider range of conditions and thereby maintain their reproductive success (Silva et al., 2019). For instance, generalist bird parasites are favoured over specialists after anthropogenic disturbances (Dharmarajan et al., 2021).

### Oncidium

We found a significant decline in pollinarium removal over time for *Oncidium* specimens across South America. This would indicate an overall decrease in pollinator availability, mainly regarding the South American oil-collecting bees *Centris* and related genera. Bee species richness, including Neotropical oil bees, is declining worldwide (Zattara and Aizen, 2021). A recent study investigated pollination rates in the Andean rainforest and found that very few visitors were effective pollinators of orchid species and that pollination limitation was occurring in all the studied populations, including an *Oncidium* species (Reyes et al., 2021). Additionally, agricultural conversion sharply reduces both the abundance and the composition of orchid bees, resulting in reduced pollination efficiency for orchid species dependent on these bees, as it was shown using long-term data from the Brazilian Amazon (Brown et al., 2024). The overall health of oil-collecting Centridini bee populations from South America lacks monitoring (Velez et al. 2017), which highlights the knowledge gap and necessity of studying these vulnerable populations.

The reproductive success of *Oncidium* species is generally low, as species often experience low visitation and fruit set, partly due to specialised pollinator requirements and the rarity of successful pollinator visits (Castro and Singer, 2019). Only a few strong enough pollinators can trigger the push and pull mechanisms of *Oncidium* flowers to successfully remove the pollinium (Thielen et al., 2021). As a result, if pollinarium removal declines over time, so will the reproductive success of these already threatened orchids, as most *Oncidium* species are self-incompatible (Castro and Singer, 2019). Our analysis revealed a significant effect of pollination strategy on pollinarium removal over time. *Oncidium* with deceiving strategies exhibited a significant decline in pollinarium removal, and the trend for species with rewarding strategies remained stable. These findings support the already established knowledge that orchids using deceiving strategies, like food deception in *Oncidium*, suffer from lower pollinator visits compared to rewarding species (Ackerman et al., 2023). Because of the ongoing insect decline, highly dependent species could be at risk of extinction.

### Ophrys

*Ophrys* orchids are deceptive, with most species relying on highly specialised pollination via sexual deception (Schiestl et al., 2000). Sexually deceptive orchids tend to exhibit higher pollinator specificity and pollination efficiency than food-deceptive orchids, which translates into higher outcrossing levels (Scopece et al., 2010). Given that Europe has experienced industrialisation and environmental changes over the last two centuries, we expected the probability of pollinarium removal of *Ophrys* species to have decreased over time (e.g. Hutchings et al., 2017). However, our findings revealed an increase in pollinarium removal over time across Europe, which could indicate that the negative effects of anthropogenic activities on natural orchid populations and pollinators occurred long before the Industrial Revolution, or that they were never impacted strongly because when pollinator populations decrease, visitation rates stay stable as the deceived male bees face a stronger pressure to find mates. Moreover, some pollinators of *Ophrys*, particularly Apidae, may have benefited from anthropogenic changes. Indeed, we found that pollinarium removal over time was significantly higher for Apidae-pollinated *Ophrys* than in species pollinated by other groups. Recent studies have shown that the distribution of a ground-nesting social bee species has widened due to more suitable environmental conditions produced by urbanisation and climate change (Gil-Tapetado et al., 2024) and nest aggregations of long-horned Eucerini bees have expanded in geographical range, supporting the idea that urbanised areas can offer suitable conditions for rare species to thrive when well maintained (Hennessy et al., 2020). In contrast, *Ophrys* species pollinated by Andrenidae exhibited a drastically different trend, with pollinarium removal declining over time. This suggests that climate change and anthropogenic activity might be negatively affecting both Andrenidae populations and their pollination services. Effectively, recent long-term studies support this trend; for instance, a study from the Netherlands shows that the proportion of insect-pollinated plant species has significantly declined over recent decades, likely driven by the reduction in pollination services (Pan et al., 2024). Furthermore, a study on the *Andrena*-pollinated *O. sphegodes* showed that the phenology of male emergence, female emergence, and flowering has been shifting over a long period, leading to the prediction that *O. sphegodes* reproductive success will be negatively affected (Hutchings et al., 2018). Interestingly, *Andrena* species seem to have restricted geographic ranges, making them especially vulnerable to environmental changes (Praz et al., 2022). It is noteworthy that most *Ophrys* species are currently assessed as non-threatened by the IUCN Red List. Even among species pollinated by Andrenidae, populations seem to remain stable despite the decreasing trend in pollinarium removal, suggesting that species might be able to compensate for reduced pollinator activity with other reproductive strategies.

## Conclusions

Although the lack of pollinator information for more than half of the *Oncidium* species and some of the *Disa* species restrained the sample size for the analysis, our results strongly suggest an association between pollinator decline and pollinator type over time. We consider a decrease in pollinarium removal over time, even marginal, as a sign of decline in pollinator availability. Our results underscore the vulnerability of specific interactions and their response to environmental changes across continents. Monitoring reproductive success in orchids offers valuable insights into the status of vulnerable, understudied pollinator groups and allows us to investigate the impact of anthropogenic pressure on specific interactions. Consequently, collecting more pollination success data from herbarium specimens could direct future conservation efforts. Further long-term studies should accommodate a larger sampling, allowing for a broader range of pollinator groups, geographical locations and species diversity.

## Acknowledgments

This research was supported by the MSc program at the Royal Botanic Gardens, Kew. CM was supported by the Dirección de Fomento de la Investigación at the Pontificia Universidad Católica del Perú (project number PI1111), the Programa Nacional de Investigación Científica y Estudios Avanzados – Prociencia (Convenio Nº 126-2020-Prociencia) and the Max Planck Partner Group. JMB and TMK were supported with a grant by the Australia-Germany Joint Research Cooperation Scheme (project number 57654754), an initiative of Universities Australia and the German Academic Exchange Service (DAAD). JMB is supported by the Australian Research Council Australian Discovery Early Career Award (project number DE220100144) funded by the Australian Government.

